# Vitamin B_12_ is neuroprotective in experimental pneumococcal meningitis through modulation of hippocampal DNA methylation

**DOI:** 10.1101/2020.01.08.898635

**Authors:** Karina Barbosa de Queiroz, Vanessa Cavalcante-Silva, Flávia Lombardi Lopes, Gifone Aguiar Rocha, Vânia D’Almeida, Roney Santos Coimbra

## Abstract

**Background:** Bacterial meningitis (BM) causes apoptotic damage to the hippocampus and homocysteine (Hcy) accumulation to neurotoxic levels in the cerebrospinal fluid of children. The Hcy pathway controls bioavailability of methyl and its homeostasis can be modulated by vitamin B_12_, cofactor of the methionine synthase enzyme. Herein, the neuroprotective potential and the underlying mode of action of vitamin B_12_ adjuvant therapy were assessed in an infant rat model of BM.

**Methods:** Eleven-day old rats were intracysternally infected with *Streptococcus pneumoniae* serotype 3, or saline, treated with B_12_ or placebo, and, 24h after infection, their hippocampi were analyzed for apoptosis in the dentate gyrus, sulfur amino acids content, global DNA methylation, transcription and proximal promoter methylation of candidate genes. Differences between groups were compared using 2-way ANOVA 2-way followed by Bonferroni post-hoc test. Correlations were tested with Spearman’s test.

**Results:** B_12_ attenuated BM-induced hippocampal apoptosis in a Hcy dependent manner (r = 0.80, *P* < 0.05). BM caused global DNA hypomethylation, however B_12_ restored this parameter. Accordingly, B_12_ increased the methylation capacity of hippocampal cells from infected animals, as inferred from the ratio S-adenosyl methionine (SAM):S-adenosyl homocysteine (SAH) in infected animals. BM upregulated selected pro-inflammatory genes, and this effect was counteracted by B_12_, which also increased methylation of CpGs at the promoter of *Ccl3* of infected animals.

**Conclusion:** Hcy is likely to play a central role in hippocampal damage in the infant rat model of BM, and B_12_ shows an anti-inflammatory and neuroprotective action through methyl-dependent epigenetic mechanisms.

## Background

The estimated incidence of bacterial meningitis (BM) is 0.7-0.9 / 100,000 people per year in developed countries and can reach values up to 10-40 / 100,000 in some African countries [1]. The most common etiologic agents of BM are *Streptococcus pneumoniae* (pneumococcus) *Neisseria meningitidis* (meningococcus) and *Haemophilus influenzae* type b (Hib). Since the advent of anti-Hib vaccine in the late 90’s, pneumococcus has become the most frequent cause of non-epidemic BM among children older than one year of age [2, 3]. Pneumococcal meningitis is associated with the highest mortality (30%) and morbidity rates in BM [4]. Between 30 and 50% of survivors develop permanent neurological sequelae, including sensory-neural deafness, cognitive deficits, and sensory-motor disabilities, seizures and cerebral palsy [5-8].

BM is characterized by intense granulocytic inflammation within the subarachnoid and ventricular spaces, extending into the perilymphatic space of the inner ear. This granulocytic inflammation results in an extensive neuronal damage, primarily in the brain cortex (CX) and hippocampus (HC). In CX, the main lesion is acute neuronal necrosis, while apoptosis is the predominant form of cell injury in HC (reviewed in [9]). HC cells undergoing apoptosis are mostly post mitotic neurons and progenitor cells distributed along the inner granule cell layer of the dentate gyrus [10, 11].

During the acute phase of BM, dramatic changes in multiple gene expression pattern may occur in CX and HC, as reported by Coimbra et al. [12]. These changes have a fundamental role regulating neurodegeneration and neuroregeneration processes, in an attempt to recovery the cognitive function in BM survivors. The HC is particularly susceptible to structural, functional and neurogenic rearrangements in response to environmental stimuli [13-15]. Several research groups have reported evidences that these alterations are regulated by epigenetic processes, defined as inheritable states of gene activity not based on changes in DNA sequence [16-18]. One of the major epigenetic mechanisms is DNA cytosine methylation [19]. Alterations in DNA methylation have been linked to human diseases, such as cancer and several neurological disorders, suggesting an important role for this epigenetic process in brain function [20]. However, the intrinsic regulatory programs and environmental factors that may modulate neuroinflammation remain unclear.

Disturbances in methyl bioavailability are associated with neurological disorders. S-adenosylmethionine (SAM or AdoMet) is the main cellular methyl donor involved in DNA, RNA, and histone methylation that may modulate gene expression via epigenetic mechanisms [21]. SAM (or AdoMet) is synthesized from ATP and methionine, which can be either dietary or formed by methylation of homocysteine (Hcy), using a methyl group of 5-methyltetrahydrofolate in a cobalamin (vitamin B_12_)-dependent reaction [22]. Methylation reactions generate S-adenosylhomocysteine (SAH or AdoHcy), a potent inhibitor of methyltransferases AdoMet-dependent. SAH (or AdoHcy) can be hydrolyzed to adenosine and Hcy; Hcy, in turn, can be remethylated by methionine synthase, or withdrawn from the methylation cycle through conversion to cysteine (Cys), in a two-step transsulfuration pathway that requires vitamin B_6_ (Figure 1). The reduction in SAM:SAH ratio is considered an indicator of decreased methylation capacity of the cells [23]; also, plasma Hcy levels can be lowered with B vitamins supplementation [24]. Our group has shown that Hcy levels in cerebrospinal fluid (CSF) are significantly higher in children with acute BM than in children with enteroviral meningitis or without infection in the central nervous system (CNS), suggesting a role of Hcy in BM pathophysiology [25]. Therefore, the hypothesis of this study is that the imbalance in the homeostasis of the sulfur amino acids could play a role in the pathophysiology of BM, and that adjuvant therapy with vitamin B_12_ might increase promoter methylation of proinflammatory genes, leading to their downregulation, thus contributing to neuroprotection of progenitor cells and post mitotic neurons in the hippocampal dentate gyrus.

**Fig. 1:**
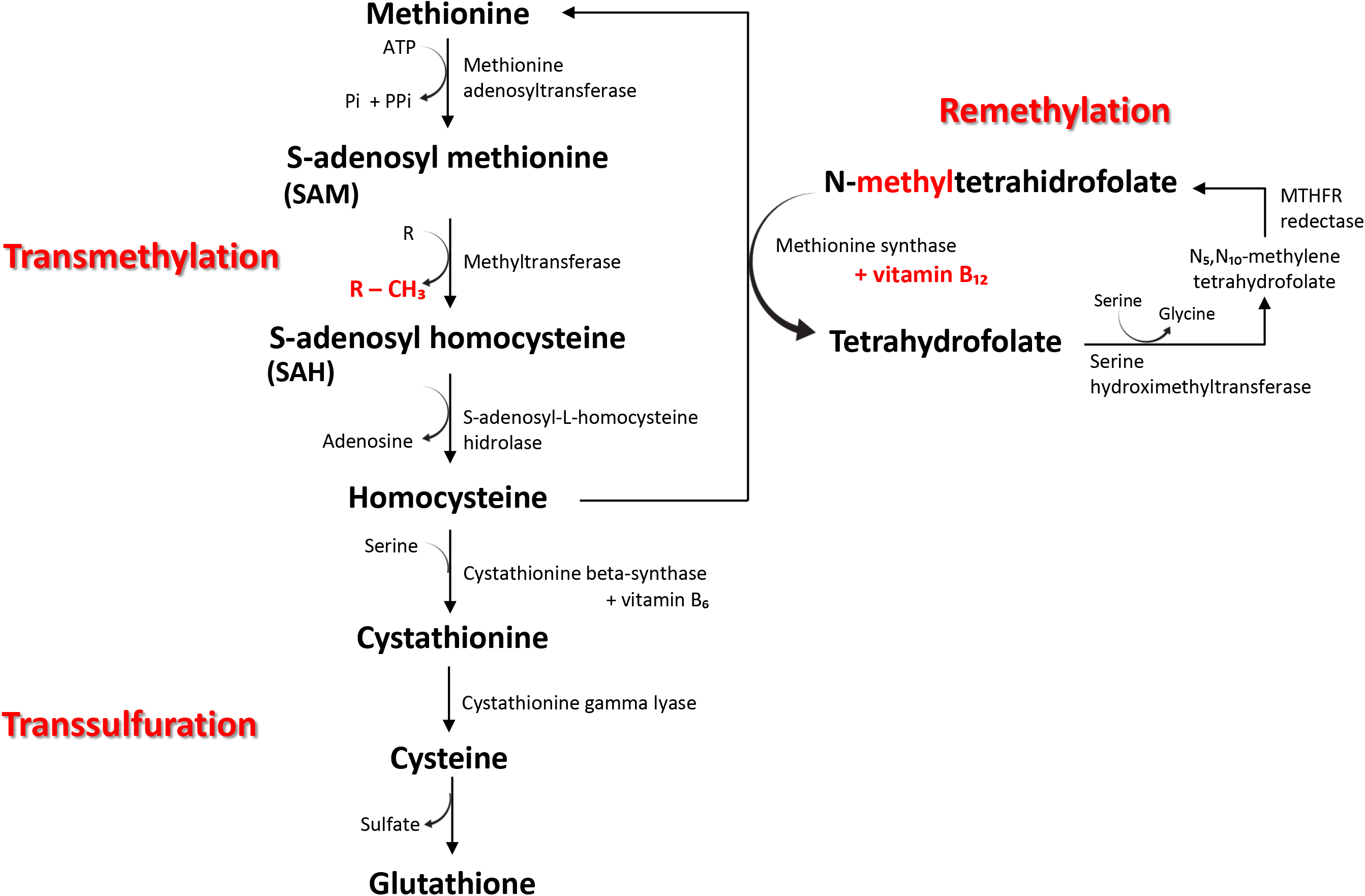
Schematic representation of homocysteine metabolism. Methionine is converted to S-adenosylmethionine (SAM) by methionine adenosyltransferase. SAM serves as a donor of methyl groups in various methylation reactions in which it is converted to S-adenosylhomocysteine (SAH). SAH is then hydrolyzed by S-adenosylhomocysteine hydrolase forming homocysteine (Hcy) and adenosine. Under normal physiological conditions, Hcy is catabolized by the transsulfuration and remethylation routes to form glutathione or methionine, respectively. Vitamin B_12_ is an essential cofactor of methionine synthase in the remethylation route. Adapted from [67].

## Methods

### Animal model and experimental design

The experiments were conducted using a well-established experimental BM model in infant rats [26]. At postnatal day 11, Wistar rats (20±2 g) were infected by intracisternal injection of 10 μL saline (0.85% NaCl) containing ~2 × 10^6^ cfu/ mL of *S. pneumoniae* (serotype 3). Animals in the control groups were sham-infected by intracisternal injection of 10 μL of sterile saline. Infected and sham-infected animals were separated into four groups according to the treatment received: 10 μL of intramuscular vitamin B_12_ (Merck, Kenilworth, NJ; 6.25 mg/ kg) (N = 10 infected - vitamin B_12_, 10 sham-infected - vitamin B_12_); or 10 μL of intramuscular saline (N = 10 infected - saline, 10 sham-infected - saline). Vitamin B_12_ or placebo was administered at three, and 18 h post-infection.

Eighteen hours after infection, all rats were weighed and clinically assessed by the following score system: 1 for comatose animals, 2 for rats that do not turn upright after positioning on the back, 3 for animals that turn within 30 s, 4 for animals that turn within less than 5 s, and 5 for rats with normal activity. Infection was documented by quantitative culture of 10 μL of CSF, and all animals were treated with 100 mg ceftriaxone /kg of body administered subcutaneously (EMS Sigma Pharma Ltda., São Paulo, Brazil) [26]. Twenty-four hours after infection, rats were euthanized by an intraperitoneal overdose of Ketamine (300 mg/ kg) + Xylazine (30 mg/ kg) (Syntec, São Paulo, Brazil).

Immediately after euthanasia, animals were perfused via the left cardiac ventricle with 7.5 mL of RNAse-free ice-cold phosphate buffered saline (PBS). Brains were removed from the skulls and the two hemispheres were divided; the right hemispheres were processed for histopathological assessment. The hippocampus from the left hemispheres were dissected in three segments, one was stored on RNA later (24 h at 4°C followed by −80°C until use), and the two others were snap frozen at −80°C to DNA and sulfur amino acids analysis.

### Brain histopathological analysis

To assess the potential of vitamin B_12_ adjuvant therapy to prevent hippocampal damage caused by BM, brains were evaluated as previously described (N = 10) [26, 27]. Briefly, the right hemisphere was fixed in 4% paraformaldehyde (Sigma-Aldrich, St. Louis, MI), embedded in paraffin, and 5 µm thick histological sections were Nissl stained with Cresyl violet. Sections were examined using a 40× objective.

Neurons of the lower blade of dentate gyrus with morphological changes characteristic of apoptosis (condensed, fragmented nuclei and/or apoptotic bodies) were counted in the largest visual field of four sections for each rat. An average score per animal was calculated from all sections evaluated, applying the following scoring system: 0–5 cells = 0, 6–20 cells = 1 and >20 cells = 2 [28].

### Quantification of sulfur amino acids in hippocampal samples

One third of each hippocampus was homogenized in PBS using a tissue homogenizer (T10 basic IKA, Staufen, Germany). For SAM and SAH measurements, protein and debris were precipitated from total homogenate tissue with HClO_4_ and centrifuged. Supernatant was then injected into a column C18 LiChroCart (5 mm, 250 mm, 64 mm). The mobile phase was applied at a flow rate of 1 mL×min^−1^ and consisted of 50 mM sodium phosphate (pH 2.8), 10 mM heptan sulfonate, and 10% acetonitrile. The UV detector had a wavelength of 254 nm. Retention time was 8.7 min for SAH and 13.6 min for SAM, a technique adapted from Blaise et al. [29]. Hcy, Cys, and glutathione (GSH) were quantified by high performance liquid chromatography (HPLC) through fluorescence detection and isocratic elution. The method developed by Pfeiffer et al. [30] was employed with slight modifications, as follows: column C18 Luna (5 mm, 150 mm, 64.6 mm), mobile phase [0.06 M sodium acetate, 0.5% acetic acid, pH 4.7 (adjusted with acetic acid), 2% methanol] and flow rate of 1.1 mL×min^−1^. The retention time was 3.6 min for Cys; 5.2 min for Hcy and 9.0 min for GSH [31]. For reduced GSH quantification, the reducing agent was not added, and the concentrations were calculated.

### Total RNA preparation

Total RNA was obtained from hippocampus using a combination of Invitrogen Trizol reagent (ThermoFisher, Waltham, MA) and chloroform (Merck) for extraction, according to the manufacturer’s protocol. Then, total RNA was purified in a column using the miRNeasy Mini Kit (Qiagen, Hilden, Germany) according to the manufacturer’s protocol. Total RNA was treated with RNase-Free DNase Set (Qiagen), following the kit’s instruction. Total RNA was quantified using the Life Technologies Qubit 2.0 Fluorometer (ThermoFisher), RNA integrity was analyzed by Agilent RNA 6000 Nano (Agilent Technologies, Waldbronn, Germany).

### Real Time quantitative Polymerase Chain Reaction for a selection of genes previously implicated in the pathophysiology of BM and DNA methylation/ demethylation

Two-hundred nanograms of total RNA were reverse transcribed into cDNA using the Applied Biosystem High Capacity cDNA RT kit (ThermoFisher). mRNA expression was quantified by real-time quantitative PCR using the Applied Biosystem Taqman system (ThermoFisher) and the ABI 7500 Fast Real-Time PCR System. Rat-specific Taqman assays were used to detect *Ccr2* [Chemokine (C-C Motif) Receptor 2] (Rn01637698_s1), *Ccl3* [Chemokine (C-C Motif) Ligand 3] (Rn01464736_g1), *Il1b* (Interleukin 1 beta) (Rn00580432_m1), *Ccl2* [Chemokine (C-C Motif) Ligand 2] (Rn00580555_m1), *Mmp9* (Matrix Metalloproteinase 9) (Rn00579162_m1), *Ocln* (Occludin) (Rn00580064_m1), *Timp1* (Tissue Inhibitor of Metallopreteinase-1) (Rn01430873_g1), *Tjp2* (Tight Junction Protein 2) (Rn01501483_m1), *Il6* (Interleukin 6) (Rn01410330_m1), *Casp3* (Caspase 3) (Rn00563902_m1), *Nfkb* (Rn01399572_m1), *Tnfa* (Tumor Necrosis Factor alpha) (Rn01525859_g1), *Il10* (Interleukin 10) (Rn01483988_g1), *Cxcl1* [Chemokine (C-X-C Motif) Ligand 1] (Rn00578225_m1), *Dnmt3a* (DNA methyltransferase 3 alpha) (Rn01027162_g1), and *Tet1* (Tet methyl cytosine dioxygenase 1) (Rn01428192_m1). Cycle thresholds (Ct) were determined based on the Taqman emission intensity during the exponential phase. Ct data were normalized by *Rplp2* (Ribosomal Protein, Large, P2) (Rn01479927_g1), and *Ppia* (Peptidylprolyl isomerase A) (Rn00690933_m1) expression, which was stably expressed in all experimental groups. The relative gene *expression* was calculated using the 2^−ΔΔCt^ method [32].

### DNA preparation

DNA of tissue samples was extracted and purified using The Wizard Genomic DNA Purification Kit (Promega, Madison, WI). Total DNA was quantified using the Life Technologies Qubit 2.0 Fluorometer (ThermoFisher) and DNA integrity was analyzed with Agilent RNA 6000 Nano (Agilent Technologies).

### Assessment of global DNA methylation

Global DNA methylation was measured using the Methyl Flash Global DNA Methylation (5mC) ELISA Easy Kit (Colorimetric) (EpiGenteK, Farmingdale, NY), according to manufacturer’s instructions. Briefly, 100 ng of genomic DNA from hippocampus (N = 6 animals per group) were bound to strip-wells specifically treated to have high DNA affinity. The methylated fraction of DNA was detected using capture and detection antibodies and then quantified colorimetrically by reading the absorbance in a micro plate spectrophotometer (450 nm). Data are presented as percentage of methylated DNA (%5mC), where the sample optical density (OD) was divided by the standard curve slope plus the OD of the DNA input.

### Assessment of DNA methylation in promoters of inflammatory genes

DNA was digested with the restriction enzyme *AluI* (20 units to 0.5 μg of DNA, 37°C, 20 h) (ThermoFisher) prior to methylation enrichment. *AluI* recognition sequence (AG^CT) does not contain the CpG dinucleotide [33].

Two-hundred and fifty nanograms of total DNA from each sample were processed with Invitrogen MethylMiner Methylated DNA Enrichment Kit (ThermoFisher), following the manufacturer’s instructions. Methylated DNA was eluted from beads using high-salt (2 M NaCl) condition. Captured DNA (methylated) and input DNA were quantified by real-time quantitative PCR using the Applied Biosystems SYBR Green system (ThermoFisher) and the ABI 7500 Fast Real-Time PCR System were used to detect the target. Rat-specific primer pairs flanking proximal promoter regions containing CpG islands and located between two *AluI* recognition sites were designed to *Ccr2* (F 5’ CAAGTGCAGTGCCTAGAGGTT 3’; R 5’ CACCTGTAATTCCAGTTTTAGGG 3’), *Ccl3* (F 5’ GGCTTCAGACACCAGAAGGA 3’; R 5’ GACTGCTGTGGTCTGCCTTAG 3’), and *Il1b* (F 5’ TCTTGGGTTGCTTGATACTGC 3’; R 5’ CATAGCCAGCCTCATGTTGA 3’). The absolute standard curve method was used to quantify the DNA amount and data were expressed as a percentage of methylated DNA [(captured DNA/ input DNA) × 100].

### Statistical analysis

Statistical analyses were performed using Graph Pad Prism version 6.0 (GraphPad Software Inc., Irvine, CA). Shapiro–Wilk test was used to verify data distribution. Differences between groups were compared using 2-way ANOVA followed by Bonferroni post-hoc test. Correlations were tested with Spearman’s test. Differences were considered statistically significant when *P* Values < 0.05.

## Results

### Meningitis and apoptosis in the inner granular layer of the dentate gyrus

All animals infected with *S. pneumoniae* had BM after 18 h p.i., as evidenced by positive bacterial titers in the CSF (~1 × 10^8^ cfu/ mL). BM decreased the activity score of the infant rats compared to sham-infected controls (*P* < 0.001). Adjuvant therapy with vitamin B_12_ did not influence bacterial titers and activity scores, but significantly reduced apoptosis in the dentate gyrus inner granular layer of infected animals, unveiling a neuroprotective effect of this potential adjuvant therapy in BM (Figure 2).

**Fig. 2:**
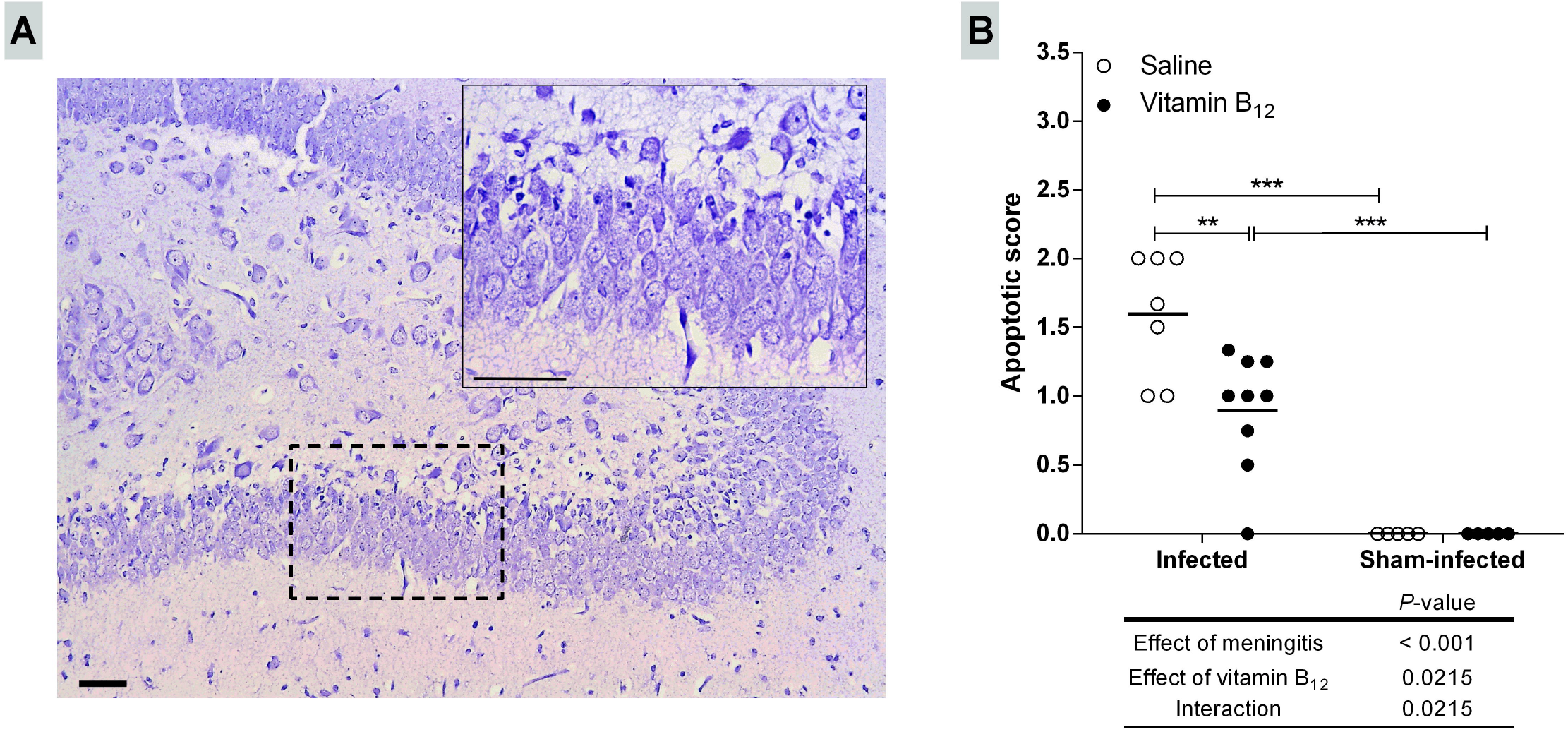
Adjuvant therapy with vitamin B_12_ attenuates apoptosis in the granular layer of the hippocampal dentate gyrus of infant rats with BM. (A) Nissl stained histological section showing the hippocampal dentate gyrus (100×) of an infected animal treated with placebo. The detail shows an amplified section of the lower blade of the granular layer (400×). Barr = 50 μm. (B) Apoptotic scores. Horizontal bars represent means. The effects of the BM and adjuvant treatment with vitamin B_12_ were compared using a 2-way ANOVA (Bonferroni post-hoc test). \*\**P* < 0.01; \*\*\**P* < 0.001.

### Quantification of sulfur amino acids in hippocampus

Hcy, Cys, total and reduced GSH were measured in the opposite hippocampi of the same animals used for histopathology. Hippocampal levels of Hcy and GSH were increased by BM. However, no effect of B_12_ was observed in these parameters (Table 1). In the infected group treated with B_12_, a positive correlation was found between hippocampal Hcy concentration and the apoptotic score in the dentate gyrus of the opposite hippocampus (r = 0.80, *P* < 0.05). In the infected group receiving placebo the apoptotic score did not correlate with Hcy concentration.

**Table 1:** Effects of BM and adjuvant therapy with vitamin B_12_ on sulfur amino acids and their metabolites in the hippocampus.

|  | Concentrations in groups (μmol/ g of protein) |  |  |  | P-values |  |  |
| --- | --- | --- | --- | --- | --- | --- | --- |
|  | Sh-Sal | Sh-B<sub>12</sub> | Infect-Sal | Infect-B<sub>12</sub> | Effect of BM | Effect of vitamin B<sub>12</sub> | Interaction |
| **Hcy** | 0.34±0.13 | 0.33±0.08 | 0.79±0.49 | 0.74±0.38 | 0.0053 | 0.8286 | 0.9079 |
| **Cys** | 1.66±0.63 | 1.8±0.46 | 2.85±1.4 | 2.13±0.67 | 0.0530 | 0.4447 | 0.2623 |
| **Total GSH** | 4.94±1.47 | 5.41±0.46 | 4.13±1.41 | 3.29±1.15* | 0.0041 | 0.6975 | 0.1698 |
| **Reduced GSH** | 3.72±0.97 | 3.98±0.52 | 2.98±0.94 | 2.57±1.05 | 0.0052 | 0.8268 | 0.3451 |

Data are expressed as means ± S.D. Statistical differences were assessed using a 2-way ANOVA (Bonferroni post-hoc test) to examine the effects of meningitis and adjuvant therapy with vitamin B<sub>12</sub>. \* denotes statistically significant differences at Bonferroni post-hoc test when compared with the respective sham-infected control (Sh-Sal or Sh-B<sub>12</sub>). Sh-Sal, sham-infected + saline; Sh-B<sub>12</sub>, sham-infected + vitamin B<sub>12</sub>; Infect-Sal, infected + saline; Infect-B<sub>12</sub>, infected + vitamin B<sub>12</sub>.
Hcy = homocysteine; Cys =Cysteine; GSH = glutathione.

To evaluate the effect of BM and of the adjuvant therapy with vitamin B_12_ on methyl bioavailability, SAM and SAH were measured in the same hippocampal samples tested for Hcy, Cys and GSH (Figure 3). SAM levels were 35% higher in infected rats treated with vitamin B_12_ as compared to infected + saline and sham-infected + B_12_ (Figure 3A). Most important, a significant increase in SAM: SAH was observed in infected rats treated with vitamin B_12_ as compared to infected + saline and sham-infected + B_12_ (Figure 3C), suggesting an increase in cell methylation capacity [34] in infected animals treated with B_12_.

**Fig. 3:**
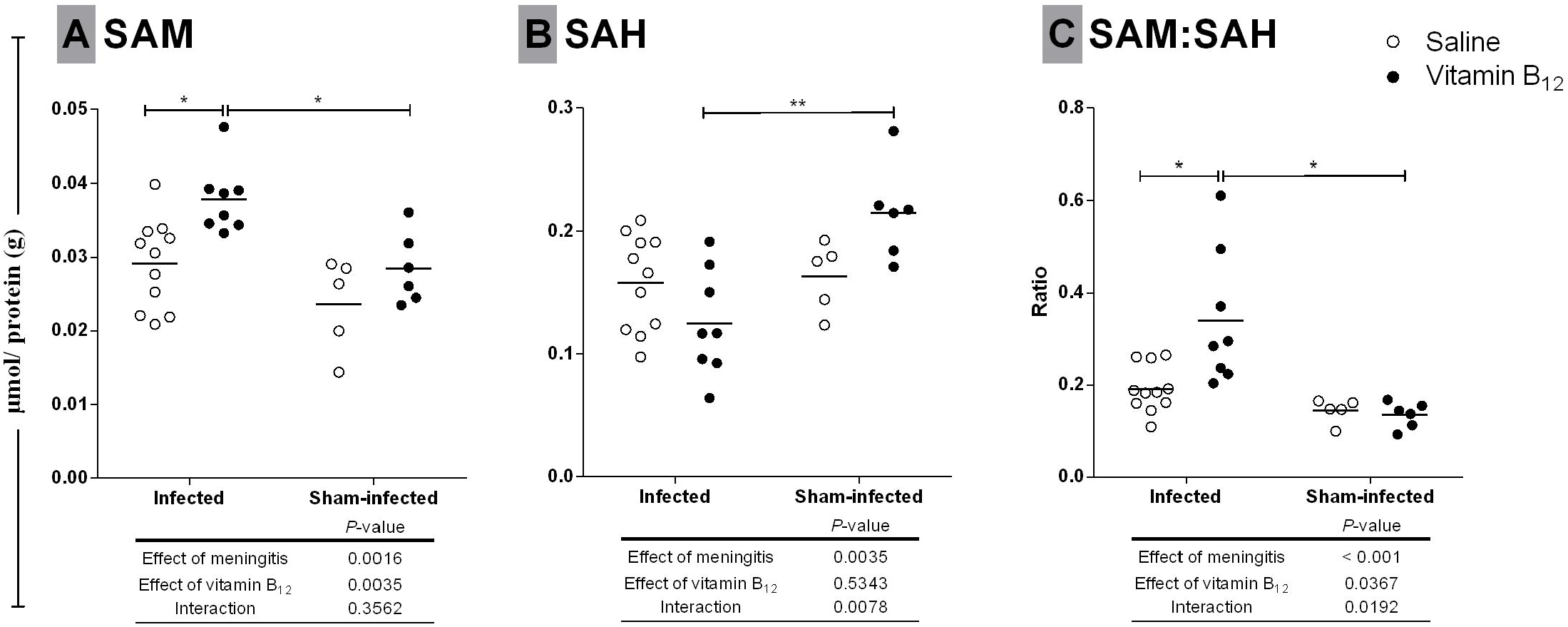
Effects of BM and adjuvant therapy with vitamin B_12_ on (A) SAM and (B) SAH concentrations and (C) SAM:SAH ratio in the hippocampus. Horizontal bars represent means. The effects of the BM and the vitamin B_12_ adjuvant treatment were compared using a 2-way ANOVA (Bonferroni post-hoc test). \**P* < 0.05; \*\**P* < 0.01. SAM = S-adenosylmethionine; SAH = S-adenosylhomocysteine.

### Percentage of global DNA methylation

To evaluate the effect of BM and adjuvant therapy with vitamin B_12_ on global DNA methylation in the hippocampus, the %5mC was measured in a colorimetric assay. The percentage of global DNA methylation was reduced by BM (*P* < 0.001), while vitamin B_12_ restored this percentage in infected animals to the control levels (Figure 4).

**Fig. 4:**
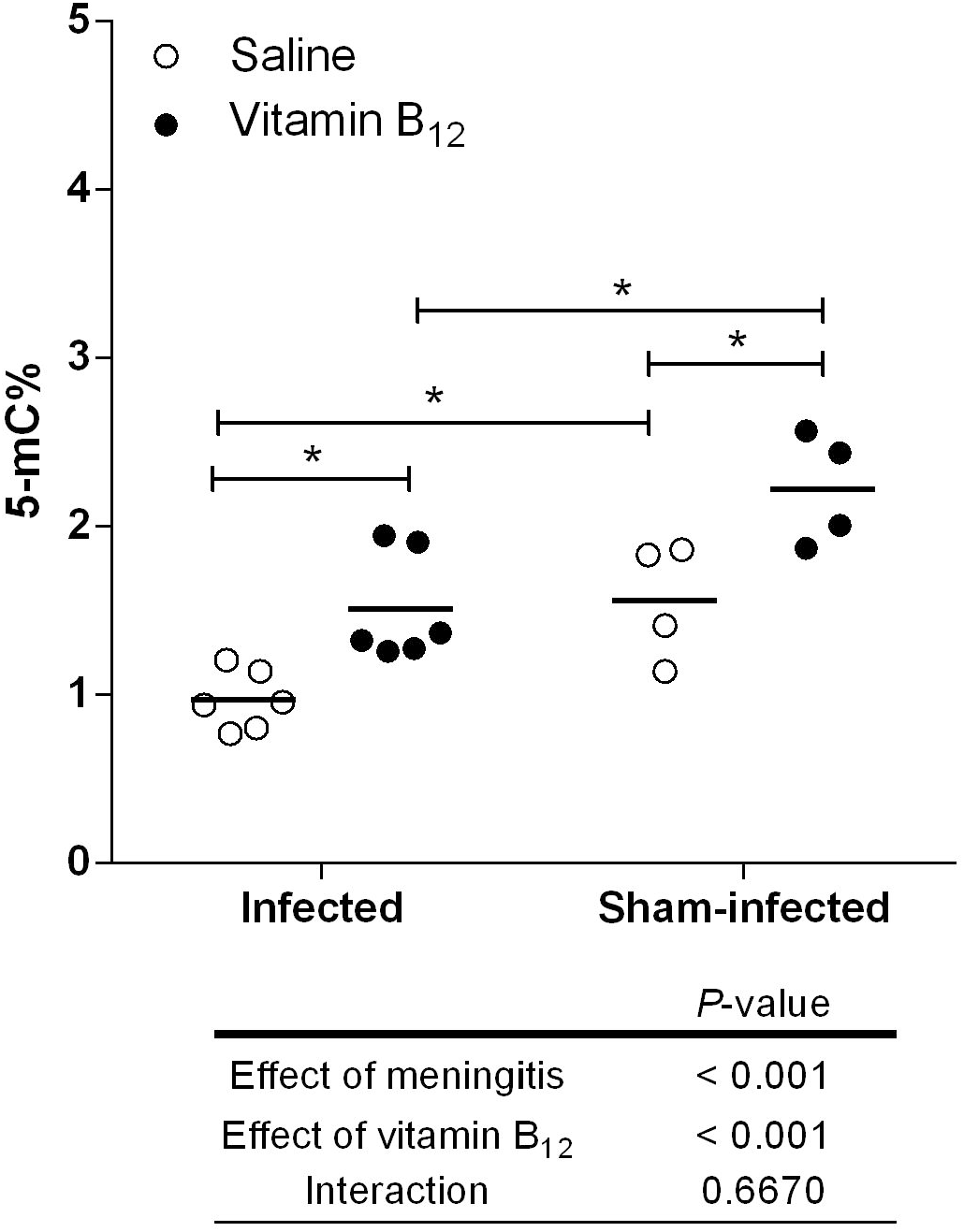
Effects of BM and adjuvant therapy with vitamin B_12_ on the percentage of global DNA methylation (5mC%) in the hippocampus. Horizontal bars represent means. The effects of the BM and the vitamin B_12_ adjuvant therapy were compared using a 2-way ANOVA (Bonferroni post-hoc test). \**P* < 0.05.

### Expression patterns of genes related to the pathophysiology of BM or DNA methylation/ demethylation

Next, the effects of BM and adjuvant therapy with vitamin B_12_ on the expression of selected genes previously implicated in the pathophysiology of BM [35] were assessed by Taqman RT-qPCR. *Ccr2*, *Ccl3* and *Il1b* had their expression increased by the infection; however, unlike other genes tested that were also affected by BM (*Mmp9, Tjp2, Ocln, Timp1, Ccl2, Il6, Casp3, Nfkb, Tnfa, Il10, and Cxcl1*) (Table 2), their increased expression was attenuated by adjuvant B_12_ (Figure 5).

**Table 2:** Effects of BM and adjuvant therapy with vitamin B_12_ on gene expression in the hippocampus.

| Genes | Relative expression in groups |  |  |  | P-values |  |  |
| --- | --- | --- | --- | --- | --- | --- | --- |
|  | Sh-Sal | Sh-B<sub>12</sub> | Infect-Sal | Infect-B<sub>12</sub> | Effect of meningitis | Effect of vitamin B<sub>12</sub> | Interaction |
| *Mmp9* | 0.0039±0.002 | 0.0061±0.003 | 0.0424±0.014 | 0.0409±0.0409 | < 0.001 | 0.9684 | 0.8385 |
| *Tjp2* | 0.0368±0.005 | 0.0737±0.033 | 0.1025±0.028* | 0.0926±0.012 | 0.0017 | 0.2419 | 0.0517 |
| *Ocln* | 0.0250±0.008 | 0.0271±0.008 | 0.0166±0.006 | 0.0120±0.005* | < 0.001 | 0.6498 | 0.2491 |
| *Timp1* | 0.0272±0.009 | 0.0139±0.004 | 1.251±0.791* | 0.9384±0.441* | < 0.001 | 0.4247 | 0.4628 |
| *Ccl2* | 0.0019±0.09 | 0.0027±0.002 | 2.165±0.344* | 2.163±0.439* | < 0.001 | 0.7075 | 0.7075 |
| *Il-6* | 0.0002±0.0001 | 0.0002±0.0001 | 0.0513±0.012* | 0.0337±0.003 | < 0.001 | 0.4040 | 0.4039 |
| *Casp3* | 0.1472±0.070 | 0.1619±0.063 | 0.0828±0.034 | 0.0991±0.042 | 0.0068 | 0.4709 | 0.9691 |
| *Nfκβ* | 0.0711±0.035 | 0.0436±0.013 | 0.1122±0.059 | 0.1712±0.104* | 0.0045 | 0.5621 | 0.1200 |
| *Tnfa* | 0.0019±0.001 | 0.0014±0.001 | 0.0355±0.017* | 0.0345±0.009* | < 0.001 | 0.8554 | 0.9583 |
| *Il-10* | 0.0001±0.001 | 0.00004±0.00004 | 0.0084±0.010 | 0.0029±0.003 | 0.0125 | 0.1941 | 0.1989 |
| *Cxcl1* | 0.0003±0.0001 | 0.0003±0.0001 | 0.4954±0.148* | 0.2427±0.051 | 0.0017 | 0.2373 | 0.2371 |

Data are expressed as means ± S.D. Statistical differences were assessed using a 2-way ANOVA (Bonferroni post-hoc test) to examine the effects of meningitis and adjuvant therapy with vitamin B<sub>12</sub>. \* denotes statistically significant differences at Bonferroni post-hoc test when compared with the respective sham-infected control (Sh-Sal or Sh-B<sub>12</sub>). Sh-Sal, sham-infected + saline; Sh-B<sub>12</sub>, sham-infected + vitamin B<sub>12</sub>; Infect-Sal, infected + saline; Infect-B<sub>12</sub>, infected + vitamin B<sub>12</sub>.

**Fig. 5:**
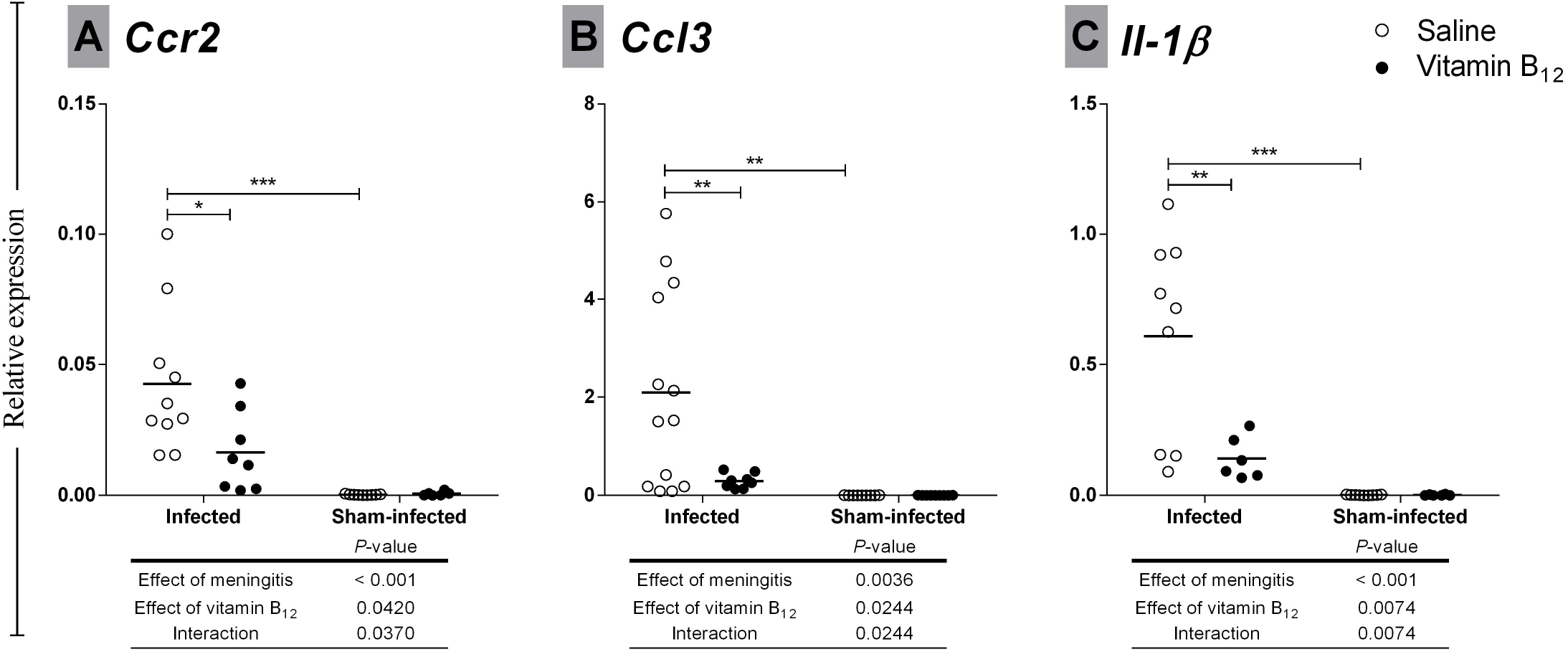
Effects of BM and adjuvant therapy with vitamin B_12_ on hippocampal gene expression. *Ccr2, Ccl3* and *Il1b* mRNA levels in hippocampus evaluated by the 2^−ΔΔCt^ method. *Rplp2* was used as a reference gene. The effects of BM and vitamin B_12_ were compared using a 2-way ANOVA (Bonferroni post-hoc test). \**P* < 0.05; \*\**P* < 0.01; \*\*\**P* < 0.001.

In the infected group treated with B_12_, Hcy concentration strongly and negatively correlated with *Ccr2* mRNA levels (r = −1.00, *P* < 0.05). In the infected group receiving placebo, SAM:SAH ratio inversely correlated with *Ccr2* expression levels (r = −0.68, *P* < 0.05), but this correlation was not found in the infected group treated with B_12_.

Transcription of DNA methyltransferase 3 (*Dnmt3a*) was influenced by the adjuvant therapy (*P* < 0.01), decreasing 1.6-fold in the infected group treated with vitamin B_12_ when compared to infected-placebo group (Figure 6A). No transcriptional change was observed for the DNA demethylase *Tet1* (Figure 6B).

**Fig. 6:**
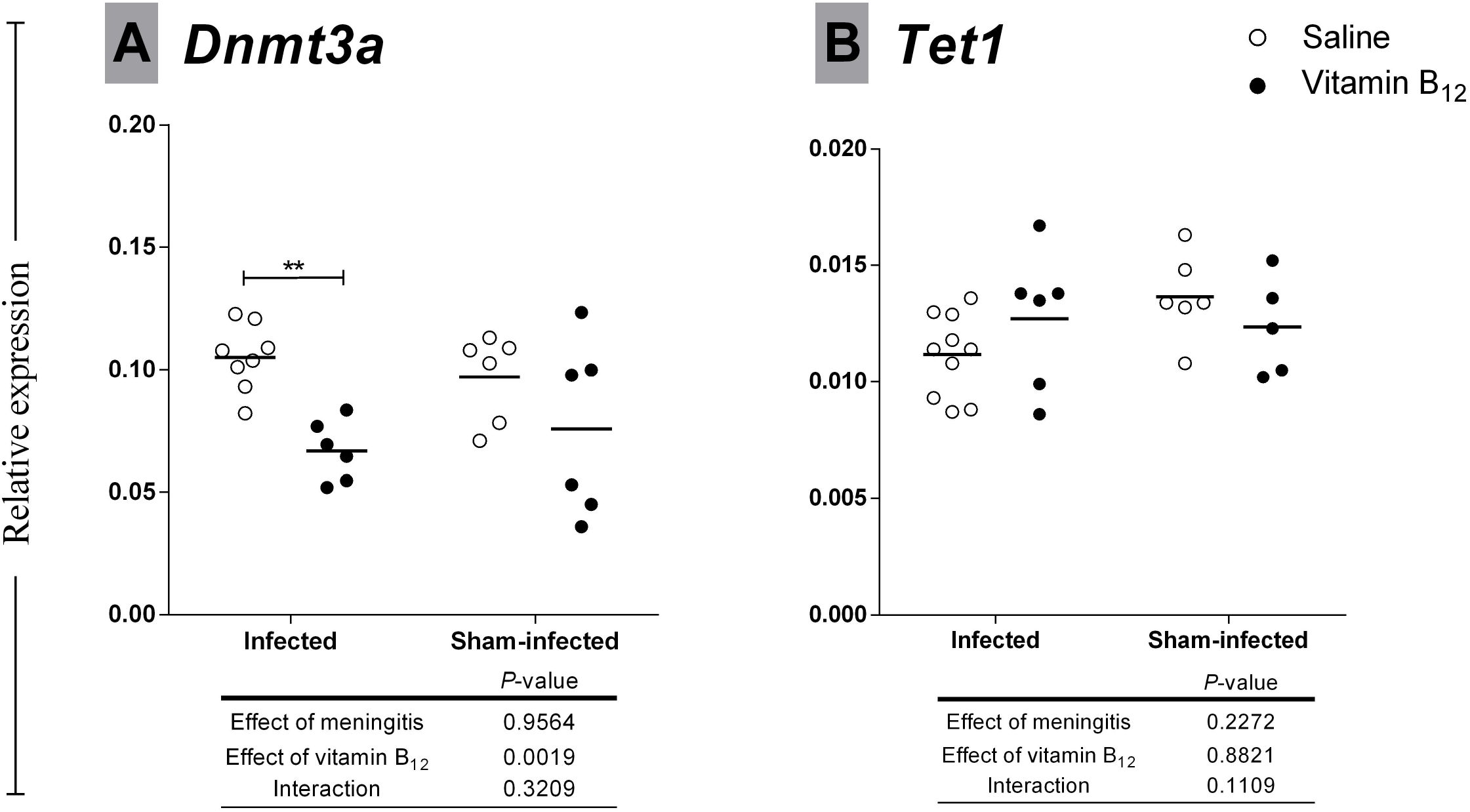
Effects of BM and adjuvant therapy with vitamin B_12_ on *Dnmt3a and Tet1* expression in the hippocampus. *Dnmt3a and Tet1* mRNA levels in hippocampus were assessed by the 2^−ΔΔCt^ method. *Ppia* was used as a reference gene. The effects of the BM and vitamin B_12_ were compared using a 2-way ANOVA (Bonferroni post-hoc test). \**P* < 0.05; \*\**P* < 0.01; \*\*\**P* < 0.001.

### Methylation of pro-inflammatory genes modulated by BM and vitamin B_12_ (<u>Ccr2</u>, <u>Ccl3</u> and <u>Il1b</u>)

To assess whether gene expression during BM was regulated by DNA methylation, differentially methylated regions (DMR) were searched, using a methyl-binding domain based enrichment approach [37], in the promoter of pro-inflammatory genes *Ccr2, Ccl3* and *Il1b*, which were upregulated by BM and downregulated by adjuvant therapy with B_12_. Hypermethylation of CpGs at the promoter region of *Ccl3* were detected in infected animals treated with B_12_ (Figures 5 and 7).

**Fig. 7:**
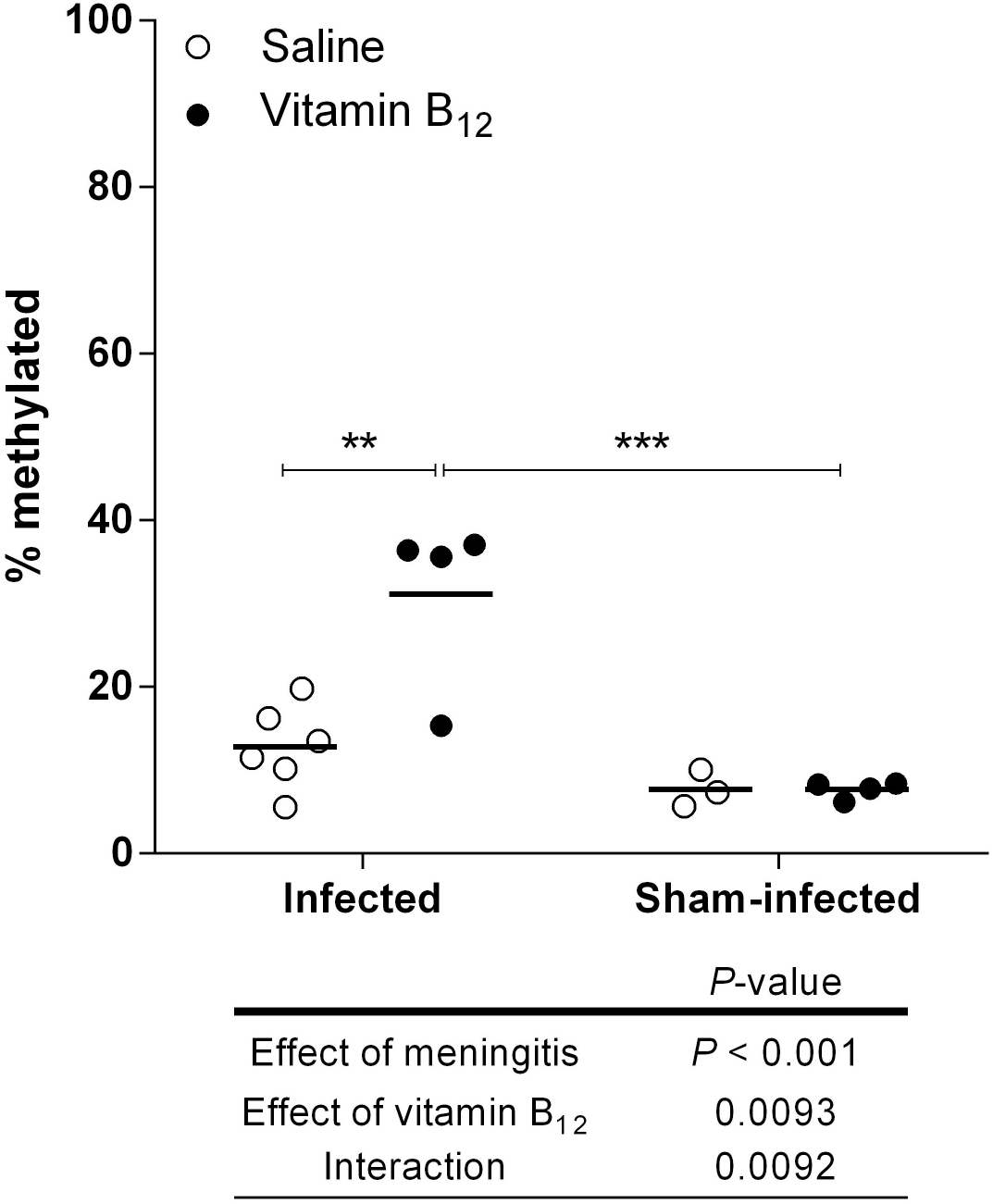
Effect of BM and adjuvant therapy with vitamin B_12_ on *Ccl3* gene methylation in the hippocampus. Percentage of methylated DNA in a region covering part of the promotor of *Ccl3* gene. The absolute standard curve method was used to quantify the DNA amount and data were expressed as a percentage of methylated DNA [(captured DNA/ input DNA) × 100]. The effects of BM and vitamin B_12_ were compared using a 2-way ANOVA (Bonferroni post-hoc test). \**P* < 0.05; \*\**P* < 0.01; \*\*\**P* < 0.001.

## Discussion

BM induces apoptosis in post-mitotic neurons distributed along the dentate gyrus inner granular layer [10, 11] and the damage to the hippocampal formation has been associated with learning and memory impairments among BM survivors [27, 38]. Previous interventional studies with B vitamin supplementation (vitamin B_6_, B_12_, and folate) demonstrated the causal relationship between vitamin B_12_ and cognitive function [39, 40]. In the present study, histomorphological analysis unveiled a significant decrease in apoptotic cell counting in the dentate gyrus inner granular layer of infected animals treated with B_12_, proving, for the first time, the neuroprotective effect of B_12_ as adjuvant therapy to pneumococcal meningitis in the infant rat model (Figure 2).

B_12_ is a cofactor of methionine synthase, which converts homocysteine into methionine modulating the sulfur amino acids pathway [22]. Our group has previously reported that Hcy concentration is increased in the CSF of children with BM compared to those with enteroviral meningitis or controls without infection in the CNS [25]. In the present study, BM increased hippocampal Hcy levels. Actually, the positive correlation found between hippocampal Hcy concentration and the apoptotic score in the dentate gyrus of infected animals treated with B_12_ supports the hypothesis that an imbalance in the sulfur amino acid homeostasis leading to Hcy accumulation may play a pivotal role in the neurodegenerative process associated to BM. Clinical investigations have shown that Hcy plasma levels are significantly higher in patients with neurologic disorders [41, 42], being a strong independent risk factor for multiple sclerosis [43], Alzheimer’s disease [44-48], and Parkinson’s disease [46, 49, 50]. Additionally, a tight correlation has been reported between the elevated Hcy plasma levels and cognitive impairment in the elderly [41, 51, 52]. Accumulated evidences show that Hcy is a potent neurotoxin, which contributes to oxidative stress in the brain [48], where it binds and activates N-methyl-d-aspartate (NMDA) receptors [53, 54], and mobilizes intracellular calcium stores [55] leading to caspase activation, DNA damage, and neuronal apoptosis *in vitro* [51, 56, 57] and *in vivo* [52, 58, 59]. Indeed, it has been reported that Hcy can upregulate *Casp3* and activate caspase 3 via p38-mitogen-activated protein kinase (p38-MAPK) [60]. Although neurotoxicity of Hcy is well documented, this is the first work to directly associate Hcy to the pathophysiology of BM, even though only correlational evidence are provided. Curiously, in the infected group receiving placebo the apoptotic score did not correlate with Hcy concentration. It is conceivable that, in this group, apoptotic scores are too high to fit within the range where correlation with Hcy concentration occurs.

During acute BM, substrate excess (Hcy) (Table 1) and increased bioavailability of the methionine synthase cofactor (supplemental B_12_) may have favored Hcy recycling, thus increasing SAM concentration, SAM:SAH ratio (Figures 1 and 3) and, consequently, rescuing the decreased percentage of global DNA methylation due to BM to normal levels (Figure 4). The biological importance of 5-methylcytosine (5mC) as a major epigenetic regulatory mechanism of gene expression has been widely recognized [19]. DNA methylation prevents genomic instability when occurring on interspersed repetitive sequences (IRSs) [61]. Actually, global DNA hypomethylation mostly reflects a decrease in DNA methylation of IRSs, which can cause genome instability [62, 63]. Methyl-deficiency due to a variety of environmental influences causes a global decrease in 5mC content (DNA hypomethylation), which has been proposed as a molecular marker in multiple pathological processes [20]. In addition, oxidative stress facilitates genome-wide hypomethylation [64]. During BM, oxidative DNA damage causes excessive activation of the DNA repair enzyme poly (ADP-ribose) polymerase (PARP). This process comes at a very high energy cost depleting NAD^+^ and ATP, thereby causing apoptosis to the granule cells of the dentate gyrus [65]. However, a link between oxidative DNA damage and global DNA methylation status in the brain during BM had not yet been proposed. The global hypomethylation reported herein in the hippocampal DNA of infected animals may further increase genome instability in the context of BM-induced oxidative stress, which could render granule cells more prone to energy collapse and apoptosis due to PARP hyper-activation. Thus, the neuroprotective effect of adjuvant B_12_ on animals with BM is likely to involve protection against oxidative DNA strand breaks.

DNA methylation also downregulates gene expression when occurring on CpG islands within promoters or other gene regulatory regions [61]. As expected, BM affected the expression of all genes tested [12]. Nevertheless, the *Ccr2*, *Ccl3* and *Il1b* expression profiles (Figure 5) were consistent with the hypothesis that treatment with vitamin B_12_ increases the bioavailability of methyl and the methylation of promoters or other regulatory regions of pro-inflammatory genes, leading to their down-regulation and, therefore, to the neuroprotection shown in Figure 2.

When Hcy and B_12_ levels are high, the enzyme methionine synthase can efficiently convert Hcy into methionine, which is then converted into SAM, the methyl donor required for DNA methylation. Accordingly, in the infected group treated with vitamin B_12_, Hcy concentration strongly and negatively correlated with *Ccr2* mRNA levels. It is noteworthy that, in the infected group receiving placebo, SAM:SAH ratio inversely correlated with *Ccr2* expression levels, but this correlation was not found in the infected group treated with B_12_. In light of these findings, it is reasonable to hypothesize that, below a given threshold, SAM:SAH ratio, which is an indicator of the methylation capacity [23], may determine the methylation level of CpG islands at some regulatory regions of *Ccr2*, down-regulating it. However, when SAM:SAH ratio is increased by adjuvant B_12_, DNA methylation in these CpG islands reaches saturation, gene expression is largely downregulated and correlation is no longer observed.

DNA methyltransferases (DNMTs) transfer the methyl group from SAM to DNA cytosine residues, resulting in a methylated DNA template and SAH, which is recycled to Hcy, as described above. The DNMT family comprises the “maintenance” methyltransferase DNMT1 (which has a preference for hemi-methylated DNA) and the “de novo” catalytic methyltransferases DNMT3A and DNMT3B [66]. Transcription of DNA methyltransferase 3 (*Dnmt3a*) was influenced by the adjuvant therapy (*P* < 0.01), decreasing 1.6-fold in the infected group treated with vitamin B_12_ when compared to infected-placebo group (Figure 6A). This result suggests a negative feedback-loop in response to the excess of substrate (SAM) in the infected rats receiving B_12_. No transcriptional change due to BM or B_12_ were observed in the DNA demethylase *Tet1* (Figure 6B), which catalyzes the conversion of 5mC into 5-hydroxymethylcytosine (5hmC) and plays a key role in active DNA demethylation [36]. Post-transcriptional or post-translational regulation of Tet1 in response to BM of adjuvant B_12_ cannot be discarded.

To assess whether gene expression during BM is regulated by DNA methylation, differentially methylated regions (DMR) were searched, using a methyl-binding domain based enrichment approach [37], at the promoter of pro-inflammatory genes *Ccr2, Ccl3* and *Il1b*, which were upregulated by BM and downregulated by adjuvant therapy with B_12_ (Figure 5). Hypermethylation of CpGs at the promoter region of *Ccl3* was detected in infected animals treated with B_12_ (Figure 7 and 8), proving the hypothesis that the mode of action of this potential neuroprotective adjuvant therapy to BM involves methyl-dependent epigenetic mechanisms of gene regulation. Unfortunately, regional DNA sequence constraints impeded the screening by qPCR of other CpG islands in the selected genes. More robust approaches such as bisulfite conversion followed by genome-wide DNA sequencing will be of great value to unveil the ensemble of genes playing a role in the pathophysiology of BM and which regulation involves DNA methylation, susceptible to be modulated by adjuvant therapy with B_12_.

**Fig. 8:**
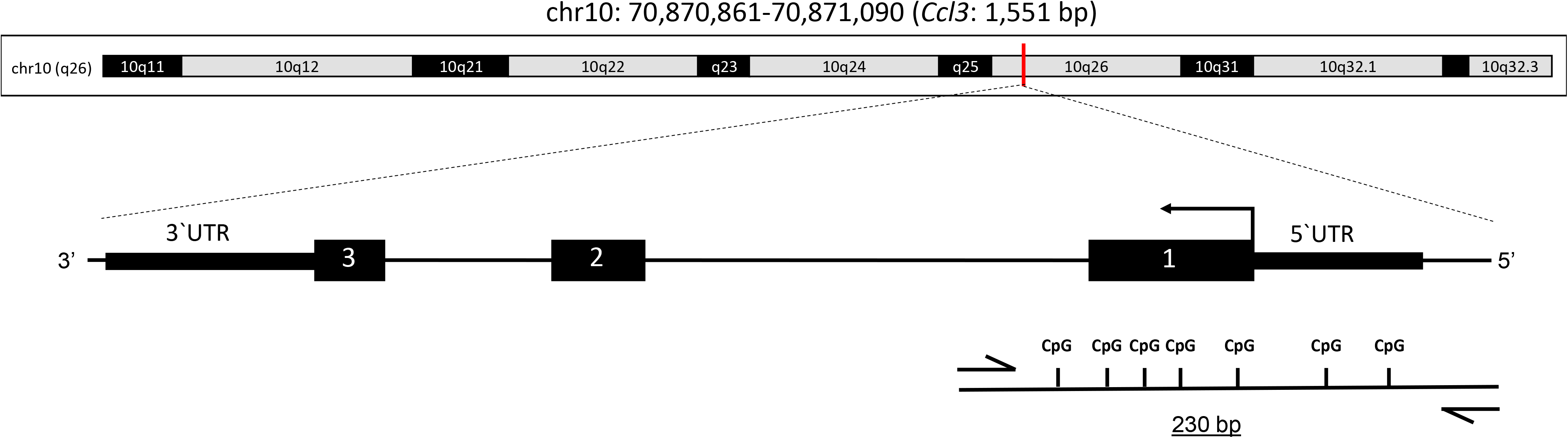
Schematic representation of *Ccl3* gene. The genome location is chromosome 10 (q26) and the gene encode a protein with 1,551 bp. Black arrow: translational start; black boxes: exons; CpG: CG dinucleotides; half arrow: primer pair chosen; dashed line: amplicon region in schematic representation.

## Conclusions

Overall, the results of this work support the hypothesis that the imbalance caused by BM in the sulfur amino acid homeostasis, which leads to Hcy accumulation in the hippocampus, may play a pivotal role in inducing apoptosis in the progenitor cells and post mitotic neurons in the hippocampal dentate gyrus. The BM-associated decrease in global DNA methylation probably contributes to genomic instability and to the previously reported over activation of PARP, a NAD-consuming DNA repair enzyme. Furthermore, adjuvant therapy with vitamin B_12_ modulates the Hcy pathway, increasing global DNA methylation, which may contribute to its neuroprotective effect on the infant rat model of BM either by reducing genome instability and by modulating the expression of critical genes to the pathophysiology of this disease.

## List of abbreviations

%5mC: percentage of methylated DNA
5hmC: 5-hydroxymethylcytosine
5mC: 5-methylcytosine
BM: Bacterial meningitis
CNS: central nervous system
CSF: cerebrospinal fluid
Ct: Cycle thresholds
CX: cortex
Cys: cysteine
DMR: differentially methylated regions
DNMTs: DNA methyltransferases
ADP-ribose: DNA repair enzyme poly
PARP: polymerase
GSH: glutathione
HC: hippocampus
Hcy: homocysteine
Hib: *Haemophilus influenzae* type b
HPLC: high performance liquid chromatography
IRSs: interspersed repetitive sequences
NMDA: N-methyl-d-aspartate
OD: optical density
PBS: phosphate buffered saline
SAH: S-adenosyl homocysteine (AdoHcy)
SAM: S-adenosyl methionine (AdoMet)

## Acknowledgements

Not applicable.

## Ethics approval

All of the experimental procedures were approved by the Ethics Committee of Care and Use of Laboratory Animals (CEUA-FIOCRUZ, protocol LW-23/17) and were conducted in accordance with the regulations described in the Committee’s Guiding Principles Manual.

## Consent for publication

Not applicable.

## Availability of data and materials

All data generated or analyzed during this study are included in this published article.

## Competing interests

The authors declared that they have no competing interests.

## Funding

This work was supported by Inova Fiocruz/Fundação Oswaldo Cruz. The funding body had no role in the design of the study, collection, analysis and interpretation of data and in writing of the manuscript.

## Author’s contributions

KBQ participated in the experiments with the animal model, analyzed and interpreted data regarding molecular biology, and contributed to the writing of the manuscript. VC-S and VD’A analyzed and interpreted the data regarding the sulfur amino acids assays. FLA participated in the conception and interpretation of the experiment to assess the methylation status of candidate genes. GAR participated in the study conception, data interpretation and manuscript writing. RSC conceived the study, analyzed and interpreted the data regarding the animal model, histology, molecular biology and sulfur amino acids assays, and wrote the manuscript. All authors critically reviewed the manuscript.

